# Hypoxia drives trastuzumab resistance through Rac1 pathway in HER2-positive breast cancer

**DOI:** 10.64898/2026.05.05.723085

**Authors:** Virginia Judith Wolos, Giulia Rocca, Marianela Abrigo, Marcela Solange Villaverde, Ezequiel Lacunza, Carla Pulero, Georgina Alexandra Cardama, Giorgio Arrigoni, Gabriel León Fiszman

## Abstract

Resistance to targeted therapy in HER2-positive breast cancer remains a clinical challenge, especially for patients with relapsed or metastatic disease. Particularly, persistent activation of hypoxia-inducible factor 1 (HIF-1) signalling is well documented in the context of trastuzumab and trastuzumab emtansine resistance. To achieve a deeper understanding of how HIF-1 activity modulates the response to anti-HER2 treatment, we functionally characterized a cellular model of hypoxia-induced drug resistance for HER2-positive breast cancer using shotgun proteomics. By global phosphoproteomics profiling, the Rac1 pathway was identified as one of the most enriched signalling networks under hypoxia. Furthermore, the selective Rac1 blockade with the 1A-116 small-molecule inhibitor sensitised HER2-positive cells to trastuzumab in both 2D and 3D culture systems. Altogether, our findings demonstrate that hypoxic conditions induce the resistance of HER2-positive breast cancer cells to targeted therapy and suggest the therapeutic potential of Rac1 inhibition to enhance trastuzumab efficacy.

**Highlights:**

- Hypoxic conditions induce trastuzumab resistance in HER2-positive breast cancer.
- Rac1 signalling was mapped under hypoxia by phosphoproteomics profiling.
- Rac1 inhibition sensitises HER2-positive cells to trastuzumab.

## INTRODUCTION

Human epidermal growth factor receptor 2 (HER2)-positive tumours comprise approximately 20 % of all human breast cancers. Of those diagnosed, over half will have complete remission at some point during treatment if HER2-targeted therapeutic monoclonal antibodies are used [1]. Trastuzumab was the first humanized monoclonal antibody developed against HER2. The binding of the antibody suppresses intracellular HER2 signalling pathways, thereby reducing cell proliferation and inducing cell cycle arrest at the G_1_ phase [2,3]. Furthermore, antibody-drug conjugates, such as trastuzumab emtansine (T-DM1), have been designed to channel the effects of cytotoxic chemotherapy. In HER2-overexpressing cells, T-DM1 causes mitotic disruption and apoptosis [4]. Despite the advances in targeting HER2, including further exploitation of antibody-drug conjugates [5], resistance to treatment remains a clinical challenge especially in cases of relapse or metastatic disease. At the molecular level, resistance to anti-HER2 therapy may occur via multiple mechanisms such as HER2 mutations, masking of HER2 epitope, HER2 heterogeneity across the tumour and activation of compensatory pathways [6].

Hypoxia-inducible factor 1 (HIF-1) is understood to be a master regulator of the cellular response to hypoxia. Like many cellular processes, the response to hypoxic stress can be harnessed by cancer cells to promote resistance to therapy [7]. HIF-1 transcription factor is a heterodimer consisting of HIF-1α and HIF-1β subunits. Its transcriptional activity is governed by the oxygen-responsive posttranslational regulation of HIF-1α. Under normal oxygen levels, this subunit is continuously synthesized and degraded by the cells. In contrast, the degradation of HIF-1α is attenuated under hypoxia, leading to the induction of HIF-1-dependent gene transcription [8]. Even so, given the high levels of HIF-1α protein expression in breast and other common human cancer tissues compared to normal tissues, the activation of HIF-1-mediated signalling in tumour cells usually occurs independently of oxygen availability. In breast cancer cells, HER2 signalling stimulates HIF-1α protein synthesis, resulting in a failure to degrade all of the HIF-1α that is expressed at normal oxygen conditions [9]. The clinical relevance of these data is underscored by the demonstration that high baseline HIF-1α expression in HER2-positive tumours is significantly associated with increased risk of metastasis and decreased patient survival [10], and with a poorer response to trastuzumab [11, 12] and T-DM1 treatments [13]. Enabled through mass spectrometry-based techniques, we functionally characterized a tumour hypoxia model for HER2-positive breast cancer. Our aim was to provide new insights into how HIF-1 activity influences the response to HER2-targeted therapy and to identify putative druggable targets for the purpose of clinical application in combined treatment approaches.

## MATERIALS AND METHODS

### Drugs

Hypoxic conditions were achieved by chemical induction with cobalt chloride (100 µM). Solid drug (CoCl_2_.6H_2_O; Cicarelli, Argentina) was dissolved in ultrapure water to a stock concentration of 25 mM, filtered and stored at -20 °C. Trastuzumab (Herceptin IV, F. Hoffmann-La Roche, Switzerland) and T-DM1 (Kadcyla, F. Hoffmann-La Roche, Switzerland) were prepared according to the manufacturer’s protocol. Both antibodies were stored at 4 °C for a maximum of three months. Based on a working concentration range from 0.001 µg/mL to 50 µg/mL, 10 µg/mL trastuzumab and 1 µg/mL T-DM1 were used for 2D cultures, while 50 µg/mL trastuzumab was applied to 3D cultures. Normal human IgG (Hemoderivados, UNC, Argentina) served as a control. The 1A-116 Rac1 inhibitor (Fig. 1) was synthesised as described by Lorenzano Menna and colleagues [14]. The compound was dissolved in ultrapure water (pH 5.5) to a stock concentration of 4.5 mM and stored at 4 °C for up to one month. Working concentrations ranged from 2.5 µM to 100 µM. Of these, 25 µM was used alone or in combination with trastuzumab across both 2D and 3D cell models.

**Fig. 1.**
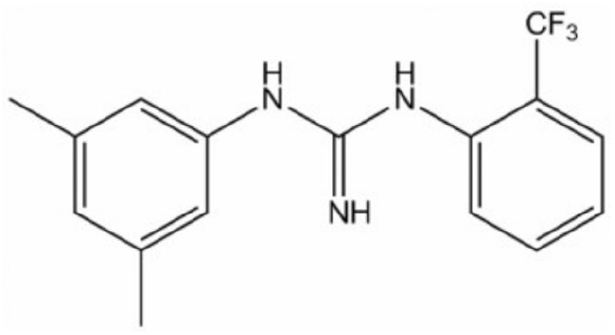
Chemical structure of the 1A-116 Rac1 inhibitor (C_16_H_16_F_3_N_3_).

### Cell culture

The BT-474 (RRID: CVCL_0179) human mammary ductal carcinoma cell line (American Type Culture Collection, USA) was cultured with Dulbecco’s Modified Eagle Medium/Nutrient Mixture F-12 (Gibco, Thermo Fisher Scientific, USA) supplemented with 10 % foetal bovine serum (Natocor, Argentina) and 40 µg/mL gentamicin, in a 5 % CO_2_/37 °C humidified incubator. Routine testing for mycoplasma was performed by PCR. Cells were harvested using trypsin-EDTA solution (Sigma-Aldrich, Merck, Germany) and seeded at 30 000 per cm^2^ for cell culture maintenance or 20 000 per cm^2^ for experiments with 2D cultures. After a 48-hour period of adherence and growth, treatments were maintained for 72 hours unless otherwise stated.

Tumour-spheroid 3D culture was conducted by applying the hanging-drop method [15]. Briefly, 20 µL drops of complete culture medium containing 10 000 cells were carefully suspended from the lid of 100 mm culture dishes. The dishes were supplemented with culture medium to prevent droplet evaporation. After a 72-hour incubation, the resulting spheroids were individually transferred to 96-well culture plates coated with 1.5 % agarose. All treatments were initiated after 48 hours of growth and maintained for 14 days by replacing half of the medium with fresh medium every two days.

### Indirect immunofluorescence

Cells seeded on coverslips were fixed with 4 % paraformaldehyde for 10 minutes. Prior to immunofluorescent staining, they were permeabilized with 0.3 % Triton X-100 at 4 °C for 10 minutes and then incubated with a blocking solution composed of 3 % BSA and 0.1 % Tween-20 for 1 hour. Incubation with anti-HIF-1α antibody (ab216842; Abcam, United Kingdom) was performed at 4 °C overnight before incubation with anti-rabbit IgG H&L Alexa Fluor 488 antibody (ab150077; Abcam, United Kingdom) in the dark for 1 hour and DAPI in the dark for 10 minutes. Finally, cells were mounted on glass microscope slides and images were captured using an Eclipse TE2000 inverted epifluorescence microscope (Nikon Instruments, Japan).

### Proteomics and phosphoproteomics analyses

#### Sample preparation

Protein extracts were obtained by homogenizing cell monolayers with RIPA buffer (20 mM Tris-HCl pH 7.5, 150 mM NaCl, 0.1 % SDS, 1 % NP-40, 0.5 % sodium deoxycholate and 1 mM EDTA) containing protease and phosphatase inhibitor cocktails (Sigma-Aldrich, Merck, Germany). Protein quantification was performed by using the Bradford method (Supelco, Merck, Germany). Then, 200 µg of each sample were lyophilized, diluted in 200 µL of washing buffer (8 M urea and 100 mM Tris-HCl pH 8.5) and loaded into a Vivacon 500 filter (Sartorius, Germany) with a molecular cut-off of 10 kDa to perform a FASP (Filter Aided Sample Preparation) protocol for protein digestion. After three washes with the indicated buffer, proteins retained in the filter were reduced with 50 mM dithiothreitol (Fluka, Honeywell, USA) in washing buffer at 55 °C for 30 minutes and alkylated with 50 mM iodoacetamide (Sigma-Aldrich, Merck, Germany) in washing buffer in the dark for 20 minutes. Then, proteins were washed twice with washing buffer, once with 100 mM ammonium bicarbonate and finally once with 50 mM ammonium bicarbonate. Protein digestion was performed by adding 100 µL of a solution containing 20 ng/µL sequencing grade modified trypsin (Promega, USA) in 50 mM ammonium bicarbonate at 37 °C overnight prior to the addition of another 25 µL of the same solution at 37 °C for 6 hours. Two extraction steps were performed with 50 mM ammonium bicarbonate to collect peptides that were finally acidified to pH 3 with formic acid, dried under vacuum and stored at-20 °C.

For the phosphoproteomics analysis, a subsequent phosphopeptide enrichment was carried out with homemade affinity chromatography microcolumns prepared by inserting 1.2 mg of titanium dioxide into C18 StageTips (Proxeon Biosystems, Denmark). Columns were conditioned twice with 50 µL of acetonitrile and twice with 50 µL of loading buffer (80 % acetonitrile and 6 % trifluoroacetic acid). Then, 190 µg of the peptide samples were dissolved in 60 µL of loading buffer and slowly loaded onto the microcolumns. The resin was washed four times with 50 µL of loading buffer and twice with 50 µL of 0.1 % trifluoroacetic acid. Finally, phosphopeptides were eluted twice with 50 µL of 15 % ammonium hydroxide (pH ˃ 10) and then twice with 50 µL of a solution containing 50 % acetonitrile and 0.1 % formic acid. Samples were dried under vacuum and stored at-20 °C.

#### Sample analyses and data processing

Liquid chromatography-mass spectrometry (LC-MS/MS) analyses were performed with an LTQ-Orbitrap XL mass spectrometer (Thermo Fisher Scientific, USA) coupled online with a nano-HPLC UltiMate 3000 system (Dionex, Thermo Fisher Scientific, USA). A label-free quantification approach was used as previously described by Arrigoni and colleagues [16]. Two technical replicates were acquired for each sample. Raw files were searched against the *Homo sapiens* section of the UniProt database version 2020-09-30 (75 093 sequences) [17] using the Andromeda search engine in MaxQuant version 1.5.1.2 [18]. Trypsin/P was used as protease, allowing up to two missed cleavages. Mass tolerances were set to 20 ppm for precursors and 0.5 Da for fragments. Additionally, match between runs was enabled with a matching time window of 0.7 minutes. Both peptide and protein FDRs were set at 1 %. The resulting protein and phosphopeptide intensities were normalized using the Cyclic-LOESS method with the web-based tool NormalyzerDE version 1.14.0 [19]. For the phosphopeptide analysis, missing values were imputed using the PhosR package version 1.14.0 [20] in R statistical software version 4.4.0 [21]. NormalyzerDE [19] was also used to assess significant differences between groups (moderated *t* test followed by Benjamini-Hochberg procedure for multiple testing correction). Proteins and phosphopeptides with adjusted p-value < 0.05 and fold change ≥ 1.2 or ≤ −1.2 were considered significantly altered.

Hierarchical clustering and heatmapping were conducted using Ward’s method with Euclidean distance measures in R [21]. The volcano plot visualization was created with the VolcaNoseR webtool [22]. The kinase-substrate relationship scoring method was applied with the PhosR package [20] in R [21]. For the functional enrichment analyses, chord diagrams were generated using the iPathwayGuide platform (AdvaitaBio Corporation, USA), while the Manhattan plot visualization was performed using the g:Profiler web server version e111_eg58_p18_f463989d [23].

### RT-qPCR

After a 6-hour exposure to 100 µM cobalt chloride, cell monolayers were homogenized in TRI Reagent RT (Molecular Research Center, USA) and RNA was purified from the lysates by extraction with chloroform. Reverse transcription reactions were performed with RevertAid Reverse Transcriptase (Thermo Scientific, Thermo Fisher Scientific, USA). Real-time quantitative PCR reactions were conducted on a CFX96 real-time PCR detection system (Bio-Rad Laboratories, USA) using TransStart Green qPCR SuperMix UDG (TransGen Biotech, China) and *VEGF* primers [24]. *GAPDH* was used as an endogenous control (forward primer: 5’-TGC ACC ACC AAC TGC TTA GC-3’, reverse primer: 5’-GGC ATG GAC TGT GGT CAT GAG-3’). Changes in gene expression were determined by the delta cycle threshold (ΔCt) method, based on the subtraction of the crossing point cycle for *GAPDH* from that of *VEGF*.

### Lactate production measurement

Lactate measurements were performed using LACT2 (Cobas, F. Hoffmann-La Roche, Switzerland). Results were normalized to the total cellular protein mass measured by the Bradford method (Supelco, Merck, Germany).

### Cell viability assays

Cell viability was assessed by crystal violet staining. The bound dye was solubilized using a mixture of ethanol and acetic acid (3:1 v/v ratio) and absorbance was measured at a wavelength of 550 nm. Alternatively, cell viability was evaluated by cell counting via the trypan blue exclusion method using a haemocytometer.

### Flow cytometry

To assess apoptosis, both attached cells harvested via trypsin-EDTA solution (Sigma-Aldrich, Merck, Germany) and detached cells were stained with FITC Annexin V Apoptosis Detection Kit II (BD Pharmingen, BD, USA). For the analysis of HER2 membrane expression, attached cells were collected as above and incubated in a 5 % CO_2_/37 °C humidified incubator for 1 hour to allow cell recovery. Then, cells were immunodecorated in the dark at 4 °C for 1 hour with anti-human HER2 APC antibody (#324408; BioLegend, USA). Samples were acquired using a BD Accuri C6 Plus flow cytometer (BD, USA) and data were analysed with FlowJo version 10.0.7 (BD, USA).

### Western blot

The HER2 and HIF-1α protein abundance was evaluated after cell treatment with 100 µM cobalt chloride for 72 hours or 25 µM 1A-116 for 24 hours, respectively. Protein extracts were obtained and quantified according to the procedure reported above for the proteomics samples. Anti-HER2 (29D8) (#2165; Cell Signaling Technology, USA), anti-β-tubulin (#2146; Cell Signaling Technology, USA), anti-HIF-1α (ab216842; Abcam, United Kingdom) and anti-β-actin (8H10D10) (#3700; Cell Signaling Technology, USA) primary antibodies were used. Immunoreactive bands were detected by electrochemiluminescence (1 M Tris base pH 8, 30 % hydrogen peroxide, 0.2 M luminol, 46 mM p-coumaric acid) and quantified using ImageJ version 1.46r [25].

### Drug combination analysis

Cell viability was assessed by crystal violet staining. The resulting data were processed by applying the Highest Single Agent synergy model in Combenefit version 2.021 [26].

### Tumour spheroid evaluation

Spheroid growth was evaluated by micrograph analysis. Image-Pro Plus version 6 (Media Cybernetics, USA) was used to measure the spheroid diameter. The spheroid volume was then calculated by using the formula for the volume of a sphere. For the histologic analysis, spheroids were collected and fixed with 4 % paraformaldehyde for 30 minutes. Then, they were dehydrated through a graded alcohol series and incubated with xylene before embedding in paraffin. Spheroid sections were cut by using a microtome, deparaffinized, rehydrated in water and finally counterstained with haematoxylin and eosin.

### Statistical analysis

All experiments were performed in at least three independent replicates. Unless otherwise stated, statistical analyses were conducted in GraphPad Prism version 9.3.1 (Dotmatics, USA). A two-sided p-value lower than 0.05 was considered statistically significant.

## RESULTS

Proteomics characterization of a cellular model of hypoxia for HER2-positive breast cancer The accumulation of the HIF-1α subunit in tumour tissue and the subsequent activation of HIF-1-mediated signalling usually occur regardless of oxygen levels [9]. Here, we assessed HIF-1α accumulation in the BT-474 HER2-positive breast cancer cell line. Remarkably, HIF-1α protein displayed a diffuse subcellular distribution in both the cytoplasm and the nucleus of the cells (Fig. 2).

**Fig. 2.**
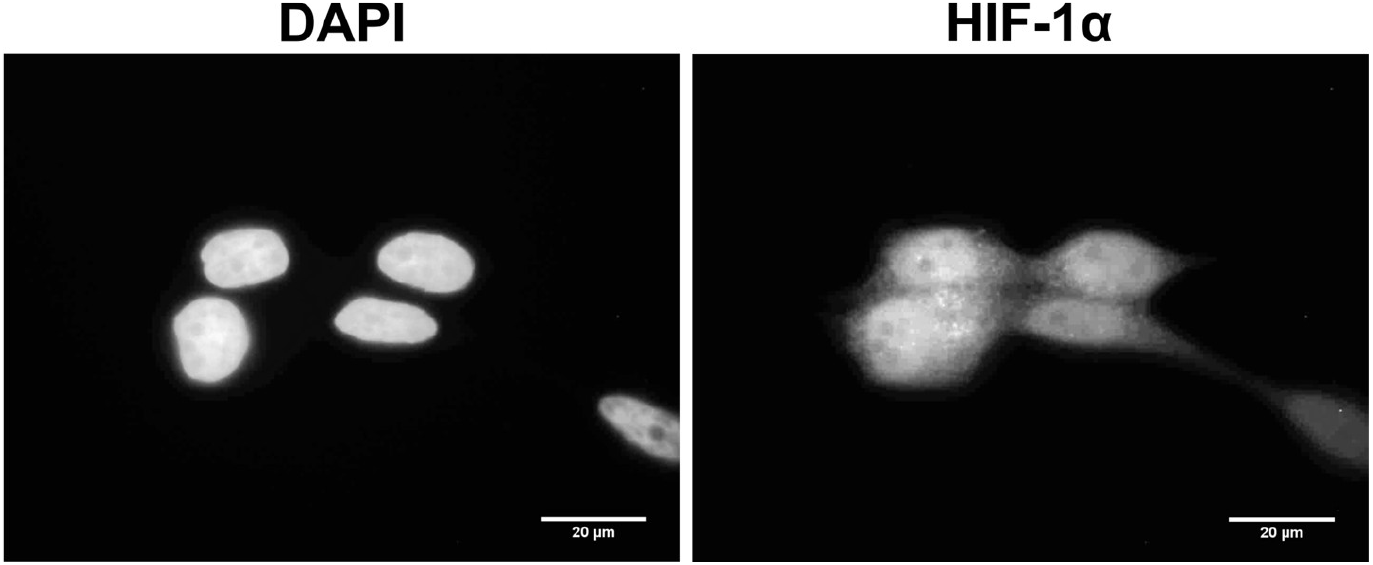
Nuclear localization of HIF-1α in HER2-positive breast cancer cells. The distribution of HIF-1α protein was examined by indirect immunofluorescence in the BT-474 cell line, showing not only a cytoplasmic but also a nuclear localization.

To study the contribution of HIF-mediated changes in the resistance to HER2-targeted therapy, hypoxic conditions were modelled on the BT-474 cell line by using cobalt chloride, which inhibits HIF-1α degradation. The hypoxic status of the cells was characterized by a proteomics analysis using a LC-MS/MS-based approach. We compared total protein extracts of cells exposed or non-exposed to cobalt chloride for 72 hours. As a result, a total of 1071 proteins were identified and 61 proteins showed statistically significant differences in their abundance between groups (Benjamini-Hochberg adjusted p-value ˂ 0.05, fold change ≥ 1.2 or ≤ −1.2) (Fig. 3a-b and Suppl. Table 1). Specifically, we found 25 downregulated and 36 upregulated proteins in hypoxic cells compared to control cells. A functional analysis using the list of 61 differentially abundant proteins led us to determine the molecular pathways enriched in the tumour hypoxia model, among which the pentose phosphate pathway, the glycolysis/gluconeogenesis and the HIF-1 signalling pathway were notable (Fig. 3c). Furthermore, we analysed upstream chemicals that may explain the pattern of differentially abundant proteins. Consistently, the results suggested oxygen deficiency as the primary upstream driver of the observed protein changes (Fig. 3d). Together, these findings indicated a hypoxia-like response in cells exposed to cobalt chloride. Increased expression of *VEGF* transcript (Fig. 3e) and elevated lactate production (Fig. 3f) further supported the LC-MS/MS results.

**Fig. 3.**
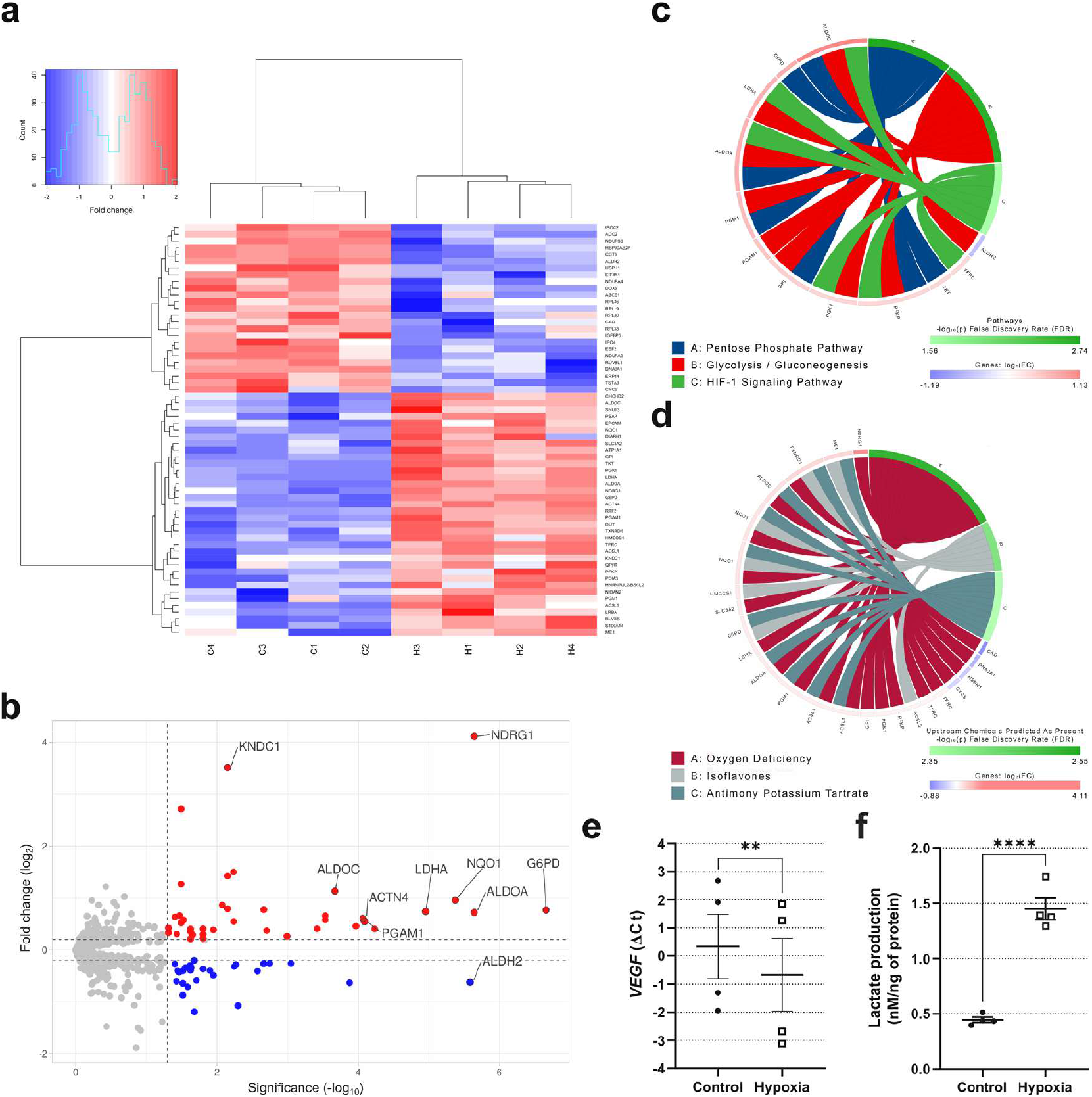
Proteomics characterization of a tumour hypoxia model for HER2-positive breast cancer. The differential protein abundance between BT-474 cells exposed (H1-H4) or non-exposed (C1-C4) to cobalt chloride was studied by LC-MS/MS (a-b). Functional analyses of the resulting 61 differentially abundant proteins highlighted the molecular pathways enriched in the tumour hypoxia model (c) and suggested oxygen deficiency as the primary upstream driver of the observed proteomics shifts (d). Increased *VEGF* expression (e) and lactate production (f) provided functional evidence of the hypoxia-like response of the cells. Data are represented as mean ± SEM; ** p < 0.01, **** p < 0.0001 by *t* test (e-f). ΔCt, delta cycle threshold.

### Hypoxia-induced cell resistance to trastuzumab and T-DM1

Using the model described above, we studied the effect of hypoxic conditions on cell response to HER2-targeted treatment with trastuzumab or T-DM1. BT-474 cells were treated with increasing concentrations of the drugs for 72 hours. A significantly higher cell viability was observed in cells treated under hypoxic conditions compared to control conditions (Fig. 4a-b). Moreover, the rate of apoptosis induced by T-DM1 was lower in hypoxic cells than in control cells (Fig. 4c). These results primarily revealed that hypoxia-mediated cell changes induced resistance to trastuzumab and T-DM1 treatments. The resistance cannot be explained by HER2 protein regulation because HER2 levels remained unaffected by hypoxic conditions (Fig. 4d-e), even when drug treatment caused a reduction in its membrane expression (Fig. 4f).

**Fig. 4.**
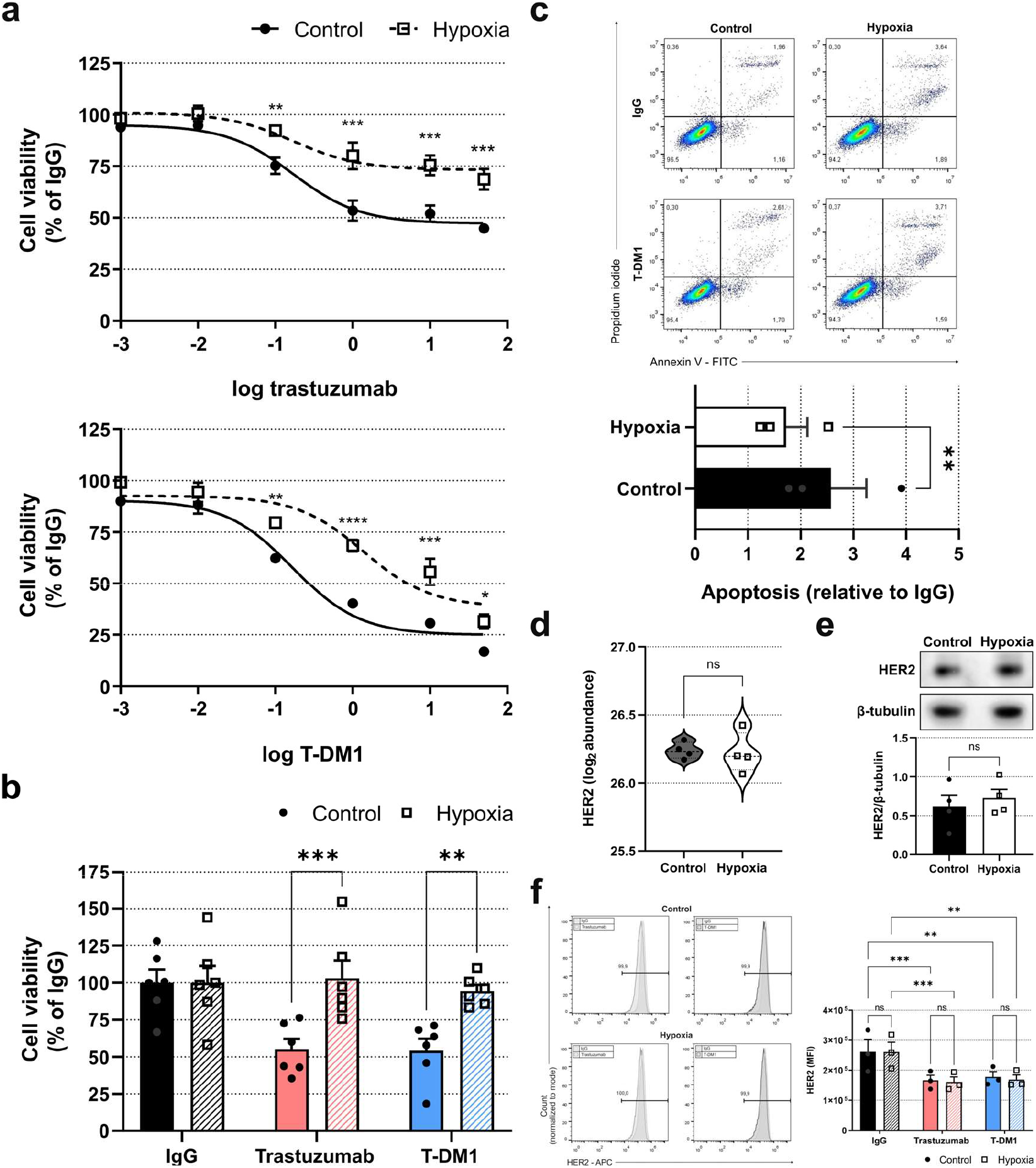
Cell resistance to HER2-targeted treatment under hypoxic conditions. Cell viability of BT-474 cells treated with trastuzumab or T-DM1 was assessed via crystal violet staining (a) and the trypan blue exclusion method (b), which exhibited cell resistance to anti-HER2 treatment under hypoxia. T-DM1-induced cell apoptosis was also attenuated by hypoxic conditions (c). However, HER2 protein abundance remained unchanged, as shown by proteomics data (d) and Western blot results (e). This was confirmed at a membrane level using flow cytometry (f). Data are represented as mean ± SEM (except for d); ns not significant, * p < 0.05, ** p < 0.01, *** p < 0.001, **** p < 0.0001 by two-way ANOVA and Bonferroni’s post-test (a-b), by *t* test (c-e) or by two-way ANOVA and Tukey’s post-test (f). MFI, mean fluorescence intensity.

### Phosphoproteomics profiling of drug-resistant cells

Mass spectrometry-based phosphoproteomics provides good insight into global cellular signalling. To pinpoint potential cellular mechanisms of hypoxia-induced resistance, we evaluated phosphopeptide-enriched protein extracts of BT-474 cells under hypoxic or control conditions using LC-MS/MS. We identified 644 phosphopeptides, 92 of which showed statistically significant differences in their abundance between groups (Benjamini-Hochberg adjusted p-value ˂ 0.05, fold change ≥ 1.2 or ≤ −1.2) (Suppl. Table 2). A kinase-substrate relationship scoring method was applied to the 92 differentially abundant phosphopeptides to identify potential kinase-substrate pairs. The analysis uncovered 17 putative protein kinases (Fig. 5a). Then, we performed a functional profiling to the list of the 17 protein kinases and their associated substrates. The Manhattan plot in Fig. 5b displays the enriched functional pathways. Beyond VEGF and ErbB2 signalling pathways, the analysis revealed the Rho GTPase signalling, particularly the Rac1 pathway.

**Fig. 5.**
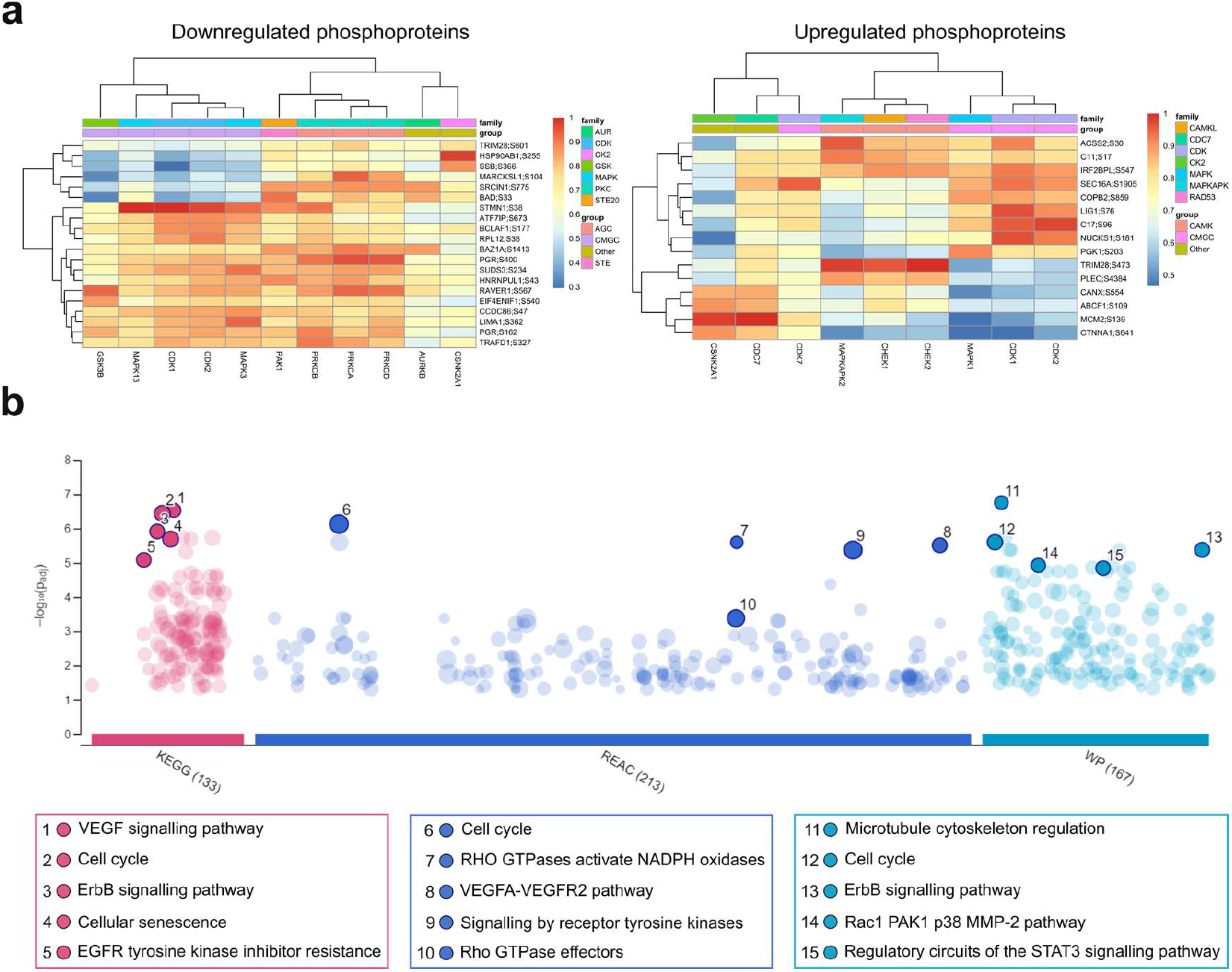
Phosphoproteomics analysis of HER2-positive breast cancer cells under hypoxia. The phosphoproteomics profile of hypoxic BT-474 cells was examined by LC-MS/MS. The application of a kinase-substrate relationship scoring method to the resulting downregulated and upregulated phosphopeptides uncovered potential kinase-substrate pairs (row dendrogram) and global relationships between kinases (column dendrogram). A higher score in the clustered heatmap denotes a better fit of a kinase to a kinase motif (a). All the kinase-substrate pairs were subjected to a functional enrichment analysis which revealed a set of statistically significantly enriched pathways in the tumour hypoxia model (b). KEGG, Kyoto Encyclopaedia of Genes and Genomes; REAC, Reactome; WP, WikiPathways.

### Improvement of trastuzumab efficacy via Rac1 inhibition

To unveil the role of Rac1 signalling pathway in hypoxia-induced resistance, we first examined whether Rac1 inhibition affected the HIF-1α protein abundance in BT-474 cells at the baseline. Remarkably, BT-474 cells treated with the specific Rac1 inhibitor 1A-116 showed significantly lower HIF-1α protein levels compared to control cells (Fig. 6a). We next evaluated the combination of trastuzumab with the 1A-116 inhibitor. BT-474 cells were treated with increasing concentrations of the drugs and cell viability was assessed after 72 hours. A drug interaction analysis using the Highest Single Agent model suggested a synergistic effect in the combined treatment (Fig. 6b). Specifically, 10 µg/mL trastuzumab plus 25 µM 1A-116 led to a statistically significant reduction in cell viability compared to single-drug treatment (Fig. 6c). Similar results were observed using a 3D culture system, which recreates the spatial cell distribution within a small tumour. BT-474 spheroids were treated with trastuzumab, 1A-116 or their combination for 14 days, and spheroid growth kinetics was assessed by calculating their volume over time. Interestingly, while trastuzumab and 1A-116 treatments inhibited spheroid growth compared to control spheroids, drug combination decreased their size (Fig. 6d). After 14 days, spheroids treated with trastuzumab plus 1A-116 showed a 88 % ± 3 % reduction in volume relative to their initial size, whereas those treated with either drug separately remained virtually unchanged (Fig. 6e). Histological sections of the spheroids showed a radial organization consisting of viable cells on the outermost layer and necrotic cells in the interior (Fig. 6f). While the necrotic core of control spheroids presented indistinct cell borders, hyperchromatic and pleomorphic nuclei, and cell detritus, spheroids treated with trastuzumab or 1A-116 showed more advanced necrotic changes such as karyorrhexis and an amorphous eosinophilic material, suggesting protein denaturation. Moreover, cells in the external layer of the spheroids treated with 1A-116 demonstrated decreased cytoplasmic eosinophilia. Of note, combined treatment caused spheroid disaggregation, precluding the histological analysis.

**Fig. 6.**
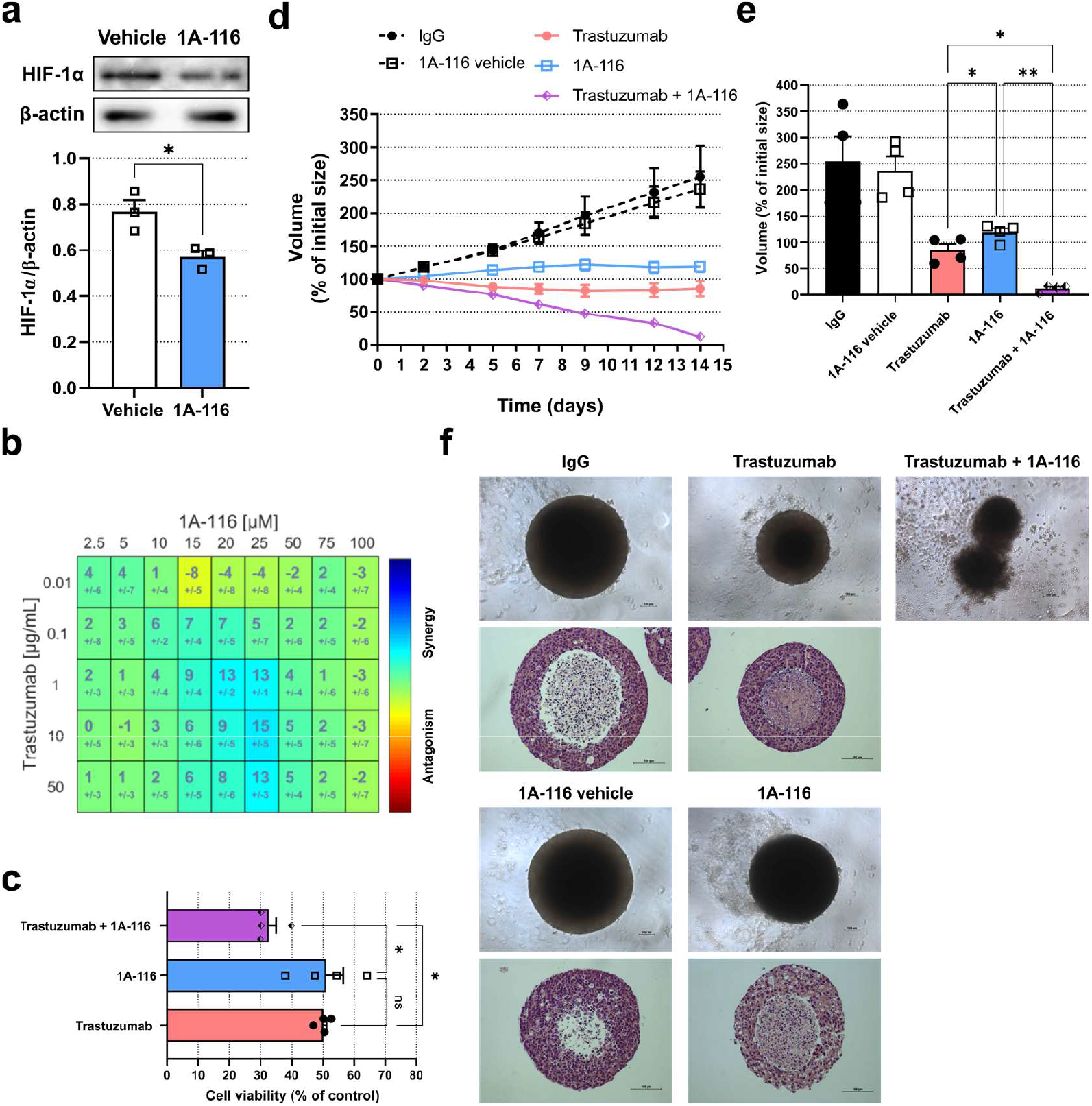
Sensitization of HER2-positive breast cancer cells to trastuzumab by Rac1 inhibition. Rac1 inhibition with the 1A-116 molecule significantly reduced HIF-1α abundance in BT-474 cells (a). The effect of trastuzumab and 1A-116 in combination was quantified and a synergy pattern was observed (b). After a 72-hour treatment, drug combination achieved a greater efficacy in reducing cell viability compared to single-drug treatment (c). In 3D cultures, while single treatments impaired spheroid growth, the combination significantly reduced the volume of the spheroids (d-e). Representative histological sections of the spheroids stained with haematoxylin and eosin are shown. Results for the combined treatment are missing due to spheroid disaggregation (f). Data are represented as mean ± SEM; ns not significant, * p < 0.05, ** p < 0.01 by *t* test (a), by two-way ANOVA and Tukey’s post-test (c) or by two-way ANOVA and Bonferroni’s post-test (e).

## DISCUSSION

The HIF-1α subunit protein abundance is frequently increased in breast cancer and other types of solid tumours [27], and it has been proposed as a potential biomarker of trastuzumab resistance [12]. Here, we demonstrated that hypoxic conditions induce the resistance of HER2-positive breast cancer cells to targeted therapy, extending available evidence for trastuzumab [11, 12, 28] and T-DM1 [13, 29]. Hypoxic conditions were achieved by using cobalt chloride in the cell culture medium. As part of this work, we conducted a functional proteomics characterization of the tumour hypoxia model using the BT-474 cell line and we confirmed a metabolic switch of the cells towards glycolysis. Resistance to treatment may lie in alternative escape pathways by which breast cancer cells circumvent HER2 inhibition. Through a mass spectrometry-based phosphoproteomics approach, the Rho GTPase effectors, particularly Rac1 pathway, were mapped under hypoxic conditions. It is known that Rac1 GTPase can be activated under multiple cellular stress conditions, including hypoxia. Increased Rac1 activity under hypoxic conditions depends on reactive oxygen species production [30]. In turn, p38 MAPK protein activation in response to hypoxia is regulated by Rac1 [31]. Indeed, Rac1 is required for the activation of stress kinases such as JNK and p38 MAPK, which play a pivotal role in DNA damage repair and cell survival [32]. Interestingly, higher levels of Rac1 and p38α MAPK were found in serum samples of metastatic breast cancer patients with HER2 overexpression [33].

Semenza and colleagues reported that HER2 signalling increases the rate of HIF-1α protein synthesis [9]. Moreover, Rac1 activity is required for the induction of the HIF-1α protein expression and transcriptional activity [31]. In this work, HIF-1α expression and its nuclear localization were detected in non-hypoxic BT-474 HER2-overexpressing cells. These basal levels of expression decreased after targeting Rac1 with the specific inhibitor 1A-116. In line with these results, recent studies have associated HIF-1α activity in tumour cells with Rac1 GTPase signalling. Rac1 activation in hepatocellular carcinoma cells under hypoxic conditions led to increased HIF-1α protein abundance and vasculogenic mimicry through Ser56 and Ser72 vimentin phosphorylation [34]. It has also been shown that Rac1 signalling drives angiogenesis via HIF-1α in triple-negative breast cancer [35]. In pancreatic cancer cells, hypoxic conditions induced gemcitabine resistance through Rac1/HIF-1α axis [36].

The Rac1 protein expression in tumours of patients with HER2-positive mammary adenocarcinoma has been associated with a poor outcome [37]. Here, we studied HER2 and Rac1 dual blockade using 2D and 3D human cell cultures. The 3D culture system reflects more accurately the complex spatial distribution of the cells within a small tumour, with a peripheral proliferation zone and a central necrotic core. In addition, our group has previously described higher HIF-1α protein expression levels in the cells surrounding the central core of BT-474 spheroids [38]. Remarkably, our results showed that Rac1 inhibition with the 1A-116 molecule significantly increased trastuzumab efficacy. Combining trastuzumab and 1A-116 led to a significant reduction in cell viability compared to single-drug treatment. Furthermore, while trastuzumab and 1A-116 single treatments inhibited spheroid growth, drug combination induced a reduction in size, leading to spheroid disaggregation. The antitumoral activity of the 1A-116 inhibitor has been studied in different types of cancer. Particularly, its preclinical efficacy and toxicology evaluation in human glioblastoma models demonstrated a strong potential for clinical translation [39].

Taken together, our findings shed new light on hypoxia-driven resistance to targeted therapy in HER2-positive human breast cancer. The Rac1 pathway emerged as one of the most enriched signalling networks under hypoxia. Rac1 inhibition with the 1A-116 molecule decreased HIF-1α abundance and increased trastuzumab efficacy, both in 2D and 3D cell cultures. Given the absence of approved therapies targeting hypoxic cells, the therapeutic potential of Rac1 inhibition holds great promise for improving the treatment of HER2-positive breast cancer with trastuzumab.

## Data statement

The datasets generated and analysed during the current study are available from the corresponding authors on reasonable request.

## Acknowledgments

We are grateful to Elisa Dora Bal de Kier Joffé and María Adela Jasnis for kindly reading and commenting on the manuscript.

## References

[1] Lester SC. The breast. In: Abbas AK, Aster JC, Kumar VK. Robbins & Cotran Pathologic basis of disease. 10 ed. Philadelphia: Elsevier; 2021. p. 1037–1064.

[2] Carter P, Presta L, Gorman CM, Ridgway JB, Henner D, Wong WL, Rowland AM, Kotts C, Carver ME, Shepard HM. Humanization of an anti-p185HER2 antibody for human cancer therapy. Proc Natl Acad Sci U S A. 1992 May 15;89(10):4285–9. doi: 10.1073/pnas.89.10.4285. PMID: 1350088; PMCID: PMC49066.

[3] Lane HA, Beuvink I, Motoyama AB, Daly JM, Neve RM, Hynes NE. ErbB2 potentiates breast tumor proliferation through modulation of p27(Kip1)-Cdk2 complex formation: receptor overexpression does not determine growth dependency. Mol Cell Biol. 2000 May;20(9):3210–23. doi: 10.1128/MCB.20.9.3210-3223.2000. PMID: 10757805; PMCID: PMC85615.

[4] Lewis Phillips GD, Li G, Dugger DL, Crocker LM, Parsons KL, Mai E, Blättler WA, Lambert JM, Chari RV, Lutz RJ, Wong WL, Jacobson FS, Koeppen H, Schwall RH, Kenkare-Mitra SR, Spencer SD, Sliwkowski MX. Targeting HER2-positive breast cancer with trastuzumab-DM1, an antibody-cytotoxic drug conjugate. Cancer Res. 2008 Nov 15;68(22):9280–90. doi: 10.1158/0008-5472.CAN-08-1776. PMID: 19010901.

[5] Ogitani Y, Aida T, Hagihara K, Yamaguchi J, Ishii C, Harada N, Soma M, Okamoto H, Oitate M, Arakawa S, Hirai T, Atsumi R, Nakada T, Hayakawa I, Abe Y, Agatsuma T. DS-8201a, A Novel HER2-Targeting ADC with a Novel DNA Topoisomerase I Inhibitor, Demonstrates a Promising Antitumor Efficacy with Differentiation from T-DM1. Clin Cancer Res. 2016 Oct 15;22(20):5097–5108. doi: 10.1158/1078-0432.CCR-15-2822. Epub 2016 Mar 29. PMID: 27026201.

[6] Swain SM, Shastry M, Hamilton E. Targeting HER2-positive breast cancer: advances and future directions. Nat Rev Drug Discov. 2023 Feb;22(2):101–126. doi: 10.1038/s41573-022-00579-0. Epub 2022 Nov 7. PMID: 36344672; PMCID: PMC9640784.

[7] Wicks EE, Semenza GL. Hypoxia-inducible factors: cancer progression and clinical translation. J Clin Invest. 2022 Jun 1;132(11):e159839. doi: 10.1172/JCI159839. PMID: 35642641; PMCID: PMC9151701.

[8] Semenza GL. Hydroxylation of HIF-1: oxygen sensing at the molecular level. Physiology (Bethesda). 2004 Aug;19:176–82. doi: 10.1152/physiol.00001.2004. PMID: 15304631.

[9] Laughner E, Taghavi P, Chiles K, Mahon PC, Semenza GL. HER2 (neu) signaling increases the rate of hypoxia-inducible factor 1alpha (HIF-1alpha) synthesis: novel mechanism for HIF-1-mediated vascular endothelial growth factor expression. Mol Cell Biol. 2001 Jun;21(12):3995–4004. doi: 10.1128/MCB.21.12.3995-4004.2001. PMID: 11359907; PMCID: PMC87062.

[10] Giatromanolaki A, Koukourakis MI, Simopoulos C, Polychronidis A, Gatter KC, Harris AL, Sivridis E. c-erbB-2 related aggressiveness in breast cancer is hypoxia inducible factor-1alpha dependent. Clin Cancer Res. 2004 Dec 1;10(23):7972–7. doi: 10.1158/1078-0432.CCR-04-1068. PMID: 15585632.

[11] Kotb RM, Ibrahim SS, Mostafa OM, Shahin NN. Potential role of CXCR4 in trastuzumab resistance in breast cancer patients. Biochim Biophys Acta Mol Basis Dis. 2022 Nov 1;1868(11):166520. doi: 10.1016/j.bbadis.2022.166520. Epub 2022 Aug 18. PMID: 35985446.

[12] Koukourakis MI, Giatromanolaki A, Boffini A, Cappelletti MR, Zanoffi L, Allevi G, Strina C, Ardine M, Milani M, Brugnoli G, Martinoffi M, Ferrero G, Bertoni R, Ferrozzi F, Harris AL, Generali D. Prospective neoadjuvant analysis of PET imaging and mechanisms of resistance to Trastuzumab shows role of HIF1 and autophagy. Br J Cancer. 2014 Apr 29;110(9):2209–16. doi: 10.1038/bjc.2014.196. Epub 2014 Apr 10. PMID: 24722179; PMCID: PMC4007245.

[13] Rediti M, Fimereli D, Mileva M, Wimana Z, Venet D, Flamen P, Guiot T, de Vries EGE, Schröder CP, Menke-van der Houven van Oordt CW, Maetens M, Majjaj S, Larsimont D, Rothé F, Sotiriou C, Gebhart G. Integrating Molecular Imaging and Transcriptomic Profiling in Advanced HER2-Positive Breast Cancer Receiving Trastuzumab Emtansine: An Analysis of the ZEPHIR Clinical Trial. Clin Cancer Res. 2025 Jan 6;31(1):110–121. doi: 10.1158/1078-0432.CCR-24-1007. PMID: 39470686.

[14] Cardama GA, Comin MJ, Hornos L, Gonzalez N, Defelipe L, Turjanski AG, Alonso DF, Gomez DE, Menna PL. Preclinical development of novel Rac1-GEF signaling inhibitors using a rational design approach in highly aggressive breast cancer cell lines. Anticancer Agents Med Chem. 2014;14(6):840–51. doi: 10.2174/18715206113136660334. PMID: 24066799; PMCID: PMC4104455.

[15] Del Duca D, Werbowetski T, Del Maestro RF. Spheroid preparation from hanging drops: characterization of a model of brain tumor invasion. J Neurooncol. 2004 May;67(3):295–303. doi: 10.1023/b:neon.0000024220.07063.70. PMID: 15164985.

[16] Aita A, Battisti I, Contran N, Furlan S, Padoan A, Franchin C, Barbaro F, Cattelan AM, Zambon CF, Plebani M, Basso D, Arrigoni G. Salivary proteomic analysis in asymptomatic and symptomatic SARS-CoV-2 infection: Innate immunity, taste perception and FABP5 proteins make the difference. Clin Chim Acta. 2022 Dec 1;537:26–37. doi: 10.1016/j.cca.2022.09.023. Epub 2022 Oct 10. PMID: 36228679; PMCID: PMC9549389.

[17] UniProt Consortium. UniProt: the Universal Protein Knowledgebase in 2025. Nucleic Acids Res. 2025 Jan 6;53(D1):D609–D617. doi: 10.1093/nar/gkae1010. PMID: 39552041; PMCID: PMC11701636.

[18] Tyanova S, Temu T, Cox J. The MaxQuant computational platform for mass spectrometry-based shotgun proteomics. Nat Protoc. 2016 Dec;11(12):2301–2319. doi: 10.1038/nprot.2016.136. Epub 2016 Oct 27. PMID: 27809316.

[19] Willforss J, Chawade A, Levander F. NormalyzerDE: Online Tool for Improved Normalization of Omics Expression Data and High-Sensitivity Differential Expression Analysis. J Proteome Res. 2019 Feb 1;18(2):732–740. doi: 10.1021/acs.jproteome.8b00523. Epub 2018 Oct 15. PMID: 30277078.

[20] Kim HJ, Kim T, Hoffman NJ, Xiao D, James DE, Humphrey SJ, Yang P. PhosR enables processing and functional analysis of phosphoproteomic data. Cell Rep. 2021 Feb 23;34(8):108771. doi: 10.1016/j.celrep.2021.108771. PMID: 33626354.

[21] R Core Team. R: A Language and Environment for Statistical Computing [software]. R Foundation for Statistical Computing, Vienna, Austria. 2021; https://www.R-project.org/. xdoi: 10.32614/R.manuals

[22] Goedhart J, Luijsterburg MS. VolcaNoseR is a web app for creating, exploring, labeling and sharing volcano plots. Sci Rep. 2020 Nov 25;10(1):20560. doi: 10.1038/s41598-020-76603-3. PMID: 33239692; PMCID: PMC7689420.

[23] Kolberg L, Raudvere U, Kuzmin I, Adler P, Vilo J, Peterson H. g:Profiler-interoperable web service for functional enrichment analysis and gene identifier mapping (2023 update). Nucleic Acids Res. 2023 Jul 5;51(W1):W207–W212. doi: 10.1093/nar/gkad347. PMID: 37144459; PMCID: PMC10320099.

[24] Medford AR, Douglas SK, Godinho SI, Uppington KM, Armstrong L, Gillespie KM, van Zyl B, Tetley TD, Ibrahim NB, Millar AB. Vascular Endothelial Growth Factor (VEGF) isoform expression and activity in human and murine lung injury. Respir Res. 2009 Apr 9;10(1):27. doi: 10.1186/1465-9921-10-27. PMID: 19358726; PMCID: PMC2674417.

[25] Schneider CA, Rasband WS, Eliceiri KW. NIH Image to ImageJ: 25 years of image analysis. Nat Methods. 2012 Jul;9(7):671–5. doi: 10.1038/nmeth.2089. PMID: 22930834; PMCID: PMC5554542.

[26] Di Veroli GY, Fornari C, Wang D, Mollard S, Bramhall JL, Richards FM, Jodrell DI. Combenefit: an interactive platform for the analysis and visualization of drug combinations. Bioinformatics. 2016 Sep 15;32(18):2866–8. doi: 10.1093/bioinformatics/btw230. Epub 2016 Apr 25. PMID: 27153664; PMCID: PMC5018366.

[27] Zhong H, De Marzo AM, Laughner E, Lim M, Hilton DA, Zagzag D, Buechler P, Isaacs WB, Semenza GL, Simons JW. Overexpression of hypoxia-inducible factor 1alpha in common human cancers and their metastases. Cancer Res. 1999 Nov 15;59(22):5830–5. PMID: 10582706.

[28] Aghazadeh S, Yazdanparast R. Activation of STAT3/HIF-1α/Hes-1 axis promotes trastuzumab resistance in HER2-overexpressing breast cancer cells via down-regulation of PTEN. Biochim Biophys Acta Gen Subj. 2017 Aug;1861(8):1970–1980. doi: 10.1016/j.bbagen.2017.05.009. Epub 2017 May 9. PMID: 28499822.

[29] Indira Chandran V, Månsson AS, Barbachowska M, Cerezo-Magaña M, Nodin B, Joshi B, Koppada N, Saad OM, Gluz O, Isaksson K, Borgquist S, Jirström K, Nabi IR, Jernström H, Belting M. Hypoxia Attenuates Trastuzumab Uptake and Trastuzumab-Emtansine (T-DM1) Cytotoxicity through Redistribution of Phosphorylated Caveolin-1. Mol Cancer Res. 2020 Apr;18(4):644–656. doi: 10.1158/1541-7786.MCR-19-0856. Epub 2020 Jan 3. PMID: 31900313.

[30] Turcotte S, Desrosiers RR, Béliveau R. HIF-1alpha mRNA and protein upregulation involves Rho GTPase expression during hypoxia in renal cell carcinoma. J Cell Sci. 2003 Jun 1;116(Pt 11):2247–60. doi: 10.1242/jcs.00427. Epub 2003 Apr 15. PMID: 12697836.

[31] Hirota K, Semenza GL. Rac1 activity is required for the activation of hypoxia-inducible factor 1. J Biol Chem. 2001 Jun 15;276(24):21166–72. doi: 10.1074/jbc.M100677200. Epub 2001 Mar 30. PMID: 11283021.

[32] Fritz G, Henninger C. Rho GTPases: Novel Players in the Regulation of the DNA Damage Response? Biomolecules. 2015 Sep 30;5(4):2417–34. doi: 10.3390/biom5042417. PMID: 26437439; PMCID: PMC4693241.

[33] Singh AK, Batra A, Upadhaya AD, Gupta S, Kp H, Dey S. Circulatory Level of Inflammatory Cytoskeleton Signaling Regime Proteins in Cancer Invasion and Metastasis. Front Oncol. 2022 Jul 7;12:851807. doi: 10.3389/fonc.2022.851807. PMID: 35875090; PMCID: PMC9300851.

[34] Zhang JG, Zhou HM, Zhang X, Mu W, Hu JN, Liu GL, Li Q. Hypoxic induction of vasculogenic mimicry in hepatocellular carcinoma: role of HIF-1 α, RhoA/ROCK and Rac1/PAK signaling. BMC Cancer. 2020 Jan 13;20(1):32. doi: 10.1186/s12885-019-6501-8. PMID: 31931758; PMCID: PMC6958789.

[35] Ren W, Liang H, Sun J, Cheng Z, Liu W, Wu Y, Shi Y, Zhou Z, Chen C. TNFAIP2 promotes HIF1α transcription and breast cancer angiogenesis by activating the Rac1-ERK-AP1 signaling axis. Cell Death Dis. 2024 Nov 13;15(11):821. doi: 10.1038/s41419-024-07223-2. PMID: 39532855; PMCID: PMC11557851.

[36] Zhu C, Hu H, Ma Y, Xiong S, Zhu D. Vav1-dependent Rac1 activation mediates hypoxia-induced gemcitabine resistance in pancreatic ductal adenocarcinoma cells through upregulation of HIF-1α expression. Cell Biol Int. 2023 Nov;47(11):1835–1842. doi: 10.1002/cbin.12074. Epub 2023 Aug 6. PMID: 37545183.

[37] Kato C, Iizuka-Ohashi M, Honda M, Konishi E, Yokota I, Boku S, Mizuta N, Morita M, Sakaguchi K, Taguchi T, Watanabe M, Naoi Y. Additional statin treatment enhances the efficacy of HER2 blockade and improves prognosis in Rac1-high/HER2-positive breast cancer. Biochim Biophys Acta Mol Basis Dis. 2024 Dec;1870(8):167458. doi: 10.1016/j.bbadis.2024.167458. Epub 2024 Aug 10. PMID: 39128642.

[38] Rodríguez CE, Reidel SI, Bal de Kier Joffé ED, Jasnis MA, Fiszman GL. Autophagy Protects from Trastuzumab-Induced Cytotoxicity in HER2 Overexpressing Breast Tumor Spheroids. PLoS One. 2015 Sep 11;10(9):e0137920. doi: 10.1371/journal.pone.0137920. PMID: 26360292; PMCID: PMC4567133.

[39] Cardama GA, Maggio J, Valdez Capuccino L, Gonzalez N, Matiller V, Ortega HH, Perez GR, Demarco IA, Spitzer E, Gomez DE, Lorenzano Menna P, Alonso DF. Preclinical Efficacy and Toxicology Evaluation of RAC1 Inhibitor 1A-116 in Human Glioblastoma Models. Cancers (Basel). 2022 Sep 30;14(19):4810. doi: 10.3390/cancers14194810. PMID: 36230732; PMCID: PMC9562863.

